# Experimental infection of specific-pathogen-free domestic lambs with *Mycoplasma ovipneumoniae* causes asymptomatic colonization of the upper airways that is resistant to antibiotic treatment

**DOI:** 10.1101/2021.05.03.442471

**Authors:** Thea Johnson, Kerri Jones, B. Tegner Jacobson, Julia Schearer, Cassie Mosdal, Steven Jones, Mark Jutila, Agnieszka Rynda-Apple, Thomas Besser, Diane Bimczok

## Abstract

*Mycoplasma ovipneumoniae* (*M. ovipneumoniae*) is a respiratory pathogen associated with the development of mild to moderate respiratory disease in domestic lambs and severe pneumonia outbreaks in wild ruminants such as bighorn sheep. However, whether *M. ovipneumoniae* by itself causes clinical respiratory disease in domestic sheep in the absence of secondary bacterial pathogens is still a matter of debate. The goal of our study was to better understand the role of *M. ovipneumoniae* as a respiratory pathogen in domestic sheep and to explore potential antibiotic treatment approaches. Therefore, we inoculated four-month-old, specific-pathogen-free lambs with field isolates of *M. ovipneumoniae* and monitored the lambs for eight weeks for colonization with the bacteria, *M. ovipneumoniae*-specific antibodies, clinical symptoms, and cellular and molecular correlates of lung inflammation. After eight weeks, lambs were treated with the macrolide antibiotic gamithromycin and observed for an additional four weeks. Stable colonization of the upper respiratory tract with *M. ovipneumoniae* was established in all four *M. ovipneumoniae*-inoculated, but in none of the four mock-infected lambs. All *M. ovipneumoniae-*infected lambs developed a robust antibody response to *M. ovipneumoniae* within 2 weeks. However, we did not observe significant clinical symptoms or evidence of lung damage or inflammation in any of the infected lambs. Interestingly, treatment with gamithromycin failed to reduce *M. ovipneumoniae* colonization. These observations indicate that, in the absence of co-factors, *M. ovipneumoniae* causes asymptomatic colonization of the upper respiratory tract of that is resistant to clearance by the host immune response as well as by gamithromycin treatment in domestic lambs.

## Introduction

*Mycoplasma ovipneumoniae* (*M. ovipneumoniae*) is a highly prevalent respiratory pathogen associated with atypical, chronic non-progressive pneumonia in sheep and goats. Colonization of the upper respiratory tract of adult sheep with *M. ovipneumoniae* is increasingly recognized across the world, with infections reported in Europe, North America, Africa, Asia, Australia and New Zealand [1–5]. In the United States, active *M. ovipneumoniae* infection was detected in 88.5% of commercial sheep flocks in a 2011 study [3]. While most cases of *M. ovipneumoniae* infection in adult sheep are thought to be asymptomatic or mild, significant production losses may occur due to reduced weight gains, lower carcass quality and increased mortality in lambs [1, 6, 7]. Moreover, *M. ovipneumoniae*-infected domestic sheep pose a significant threat to wild ruminant populations such as bighorn sheep (*Ovis canadensis*) [8, 9], Dall’s sheep (*Ovis dalli dalli*) [10], Argali sheep [11], and Norwegian Muskox (*Ovibos moschatus*) [12], where *M. ovipneumoniae* infection causes severe pneumonia outbreaks with up to 100% mortality.

However, it is still unclear to what extent *M. ovipneumoniae* causes clinical respiratory disease in sheep in the absence of other pathogens. Several studies have demonstrated that *M. ovipneumoniae* greatly increases the susceptibility for infection with other opportunistic pathogens such as *Mannheimia haemolytica* and *Bibersteinia trehalosi*, which then leads to proliferative interstitial pneumonia with severe clinical symptoms and significant mortality [9, 13–15]. However, infection studies with *M. ovipneumoniae* alone have yielded conflicting results. Thus, respiratory symptoms and lung pathology were observed in a subset of sheep following endobronchial application of *M. ovipneumoniae* in two early studies [16, 17], and more recently, Du et al. described coughing, wheezing and increased body temperatures in Bashbay lambs experimentally inoculated with *M. ovipneumoniae* [18]. In contrast, Buddle et al. [19] did not find evidence of pneumonia in colostrum-deprived lambs after experimental *M. ovipneumoniae* infection. In bighorn sheep, inoculation with cultured strains of *M. ovipneumoniae* did not result in clinical respiratory disease [8], whereas natural transmission of *M. ovipneumoniae* between animals led to severe bronchopneumonia [20]. These conflicting findings point to a need for further experimental investigations.

While the contribution of *M. ovipneumoniae* to sheep respiratory disease is widely acknowledged and elimination of *M. ovipneumoniae* colonization in domestic sheep could prevent severe disease outbreaks in bighorn sheep and other wild ruminants, no vaccines or treatments to combat the infection are currently approved. While experiments have demonstrated the *in vitro* susceptibility of most *M. ovipneumoniae* isolates to a wide range of macrolide, tetracycline and fluoroquinolone antibiotics [4, 21], no studies on antibiotic treatment of *M. ovipneumoniae* infection *in vivo* have been published.

To better understand the role of *M. ovipneumoniae* as a respiratory pathogen in domestic sheep and to explore potential antibiotic treatment approaches, we performed an *M. ovipneumoniae* infection and antibiotic treatment study in four-month-old specific-pathogen-free lambs. Intranasal inoculation of lambs with a native field isolate of *M. ovipneumoniae* resulted in consistent colonization of the upper respiratory tract and robust *M. ovipneumoniae*-specific humoral immunity. However, we did not observe clinical symptoms or evidence of lung damage or inflammation in any of the infected lambs. Interestingly, treatment with the macrolide antibiotic gamithromycin, which has been used successfully to treat clinical *Mycoplasma*-associated pneumonia in cattle, goats and pigs [22–24], failed to reduce *M. ovipneumoniae* colonization in the infected lambs. Overall, these data suggest that, in the absence of other opportunistic pathogens, *M. ovipneumoniae* behaves like an upper respiratory tract commensal that is not affected by the host antibody response or a macrolide antibiotic gamithromycin, which is commonly used to treat respiratory disease in ruminants.

## Methods

### Animals and husbandry

All animal experiments in this study were approved by the Institutional Animal Care and Use Committee (IACUC) of Montana State University, protocol #2019-95. To generate specific-pathogen-free lambs, 15 mixed-breed ewes (Rambouillet, Suffolk, Targhee, Columbia, and/or Hampshire) 4-5 years of age were purchased several weeks before the projected lambing date from a local sheep farmer. Pregnancy was confirmed using abdominal ultrasound, and *M. ovipneumoniae* exposure was determined by serology and nasal swab PCR, as described below. Prior to lambing, ewes were fed hay from the Johnson Family Livestock Facility farm, grain, and an appropriate vitamin/mineral supplement.

To derive lambs free from *M. ovipneumoniae* and other facultative respiratory pathogens, we used supervised lambing and motherless rearing, as previously described [25, 26]. Around the projected lambing date, ewes were monitored around the clock for signs of imminent delivery. Ewes developing signs of labor were separated from the flock and moved into a designated lambing pen, and lambs were manually delivered onto sterile towels placed on top of a clean plastic sheet and then were transferred into a separate, heated nursery area within the Johnson Family Livestock Facility (JFLF) ABSL-2 laboratory. All animal care personnel showered and changed into sterile PPE prior to entering the nursery, and personnel responsible for the lambs did not have any contact with other non-SPF sheep for the duration of lambing. Lambs were housed in heated animal rooms (15.5-16.8 °C) inside the JFLF in groups of 5 – 10 animals, with siblings and lambs of similar ages grouped together. During the first 24 h, lambs were bottle-fed a commercial colostrum replacer (Rescue Lamb & Kid Colostrum Replacer, Lifeline Nutrition Solutions/APC, Ankeny, IA), which contains bovine serum as an antibody source. Animals were trained to use bucket feeders and were fed a commercial lamb milk replacer diet (Hubbard Feeds, Mankato, MN) for 5 weeks, with ad libitum access to hay and water. Lambs were gradually weaned onto pelleted lamb food starting at 36-48 days of age days by decreasing the concentration of milk replacer to 50% for two weeks as previously described [25].

### Health monitoring

Approximately 2 months prior to lambing, ewes were screened for *M. ovipneumoniae, Coxiella burnetii* and *Mycobacterium avium paratuberculosis* (MAP) exposure by serum ELISA and for parainfluenza 3 (PI3) exposure by serum-based virus neutralization assay. Nasal swabs from the ewes were also analyzed for *M. ovipneumoniae* by PCR.

SPF lambs at the JFLF were screened daily for signs of illness or discomfort. In addition, SPF-status of the lambs was confirmed by testing nasal swab samples for *M. ovipneumoniae* infection by quantitative PCR at birth, at the age of 1 month, and seven days prior to the start of the experiment. At 3 months of age, nasal swabs from a subset of lambs were analyzed for *M. ovipneumoniae* by PCR and for the presence of Pasteurellaceae by conventional aerobic culture. All laboratory assays were performed at the Washington Animal Disease Diagnostic Laboratory (WADDL, accredited by the American Association of Veterinary Laboratory Diagnosticians).

Following experimental *M. ovipneumoniae* challenge or mock treatment, lamb health status was assessed twice daily by trained JFLF personnel. The following health parameters were assessed and were used to calculate a clinical disease score: (1) general behavior, (2) appetite, (3) rectal temperature, (4) respiratory symptoms, and (5) any medications administered exclusive of the experimental treatment provided at 8 weeks p.i., as detailed in **Table 1**. Daily scores represent the sum of the two values obtained for each day, and weekly scores represent the sum of all 14 values collected over a 7-day period. In addition, lamb body weights were determined on days 20, 32 and 55 post challenge.

### Experimental infection with Mycoplasma ovipneumoniae

Out of thirty live lambs born in our flock, four SPF lambs aged between 15 and 16 weeks were selected for experimental infection with *M. ovipneumoniae*, and four additional lambs, matched for age and sex, were selected as a control group. Animal details are listed in **Table 2**. The SPF lambs were infected using pooled nasal wash fluids from lambs previously determined to be infected with *M. ovipneumoniae*, following published protocols [20, 27, 28]. Animals housed at MSU’s Red Bluff Research Ranch, aged 11-13 weeks, were used as donor lambs for nasal washes. We selected eight lambs (4 ewes and 4 wethers) with mild respiratory disease symptoms that were confirmed to be infected with *M. ovipneumoniae* based on PCR analysis of nasal swabs. To obtain nasal wash fluids, the *M. ovipneumoniae*-infected lambs were restrained using halters, with the heads kept in a slightly lowered position. Nasal washes were performed by squirting 2 x 15 mL of sterile PBS into each nostril and collecting the flush fluid into a clean polyethylene bag. Nasal wash fluids were transferred to the laboratory and were immediately pooled, diluted 1: 1 with tris-buffered saline, and then were treated with ceftiofur (100 μg/mL, MWI Animal Health, Boise, ID) for 2 h at 37°C to reduce contamination with *Pasteurellacea*. Microbiological analysis of the pooled nasal wash fluid confirmed that both *Mannheimia haemolytica* and *Bibersteinia trehalosi* were present before ceftiofur treatment, but were undetectable after the treatment. We then inoculated sheep with 50 mL of the treated nasal wash fluid (*M. ovipneumoniae* group) or PBS (control group) by infusing 15 mL into each nostril, 10 mL into the oral cavity and 5 mL into each conjunctival sac. Following inoculation, the *M. ovipneumoniae* group and the control group were housed in separate rooms of the JFLF to prevent aerosol transmission of *M. ovipneumoniae*. Each room had its own set of equipment so that no equipment was shared between the rooms. For the duration of the infection experiment, staff first performed all necessary husbandry procedures on the control animals before entering the room with the infected animals and showered and changed immediately after leaving the room with the infected lambs.

### Antibiotic treatment

Eight weeks after experimental challenge with *M. ovipneumoniae* or mock challenge with PBS, all lambs were treated with 6 mg/kg BW gamithromycin (Zactran®, Boehringer Ingelheim, Ridgefield, CT) two times over 5 days by subcutaneous injection.

### Collection and analysis of bronchoalveolar lavage fluids

To collect bronchoalveolar lavage fluid (BAL), a flexible fiber-optic endoscope measuring 6.6 mm x 100 cm (VFS-2B VetVu, Swiss Precision Products, Inc, Oxford, MA) was introduced into the trachea under local anesthesia with lidocaine. Lavage was performed by instilling 60 mL of sterile saline into the lungs and then aspirating the liquid again; approximately 20 mL of fluid were routinely recovered from the sheep. BAL fluid was stored on ice until transfer to the laboratory, where cells were harvested by centrifugation at 400 *g* for 10 min. Supernatants were then analyzed for evidence of lung damage using the Lactate Dehydrogenase Assay Kit (Abcam, Cambridge, UK). Cell pellets were processed for absolute cell counts using a hemocytometer and for differential cell counts using cytospin preparations stained with a DippKwik stain (ThermoFisher, Waltham, MA). At least 300 cells from each sample were classified by a scientist blinded to the treatment of the animals.

### *Detection of* Mycoplasma ovipneumoniae *infection*

PCR detection of *M. ovipneumoniae* infection was performed at the Washington Animal Disease Diagnostic Laboratory (WADDL, Pullman, WA) using nasal swab samples and standard protocols [29]. Briefly, probe based quantitative PCRs were performed with the following primers and probe: forward: 59-GGG GTG CGC AAC ATT AGT TA-39; reverse: 59-CTT ACT GCT GCC TCC CGT AG-39; and probe: 59-6-FAM-TTA GCG GGG CCA AGA GGC TGT A-BHQ-1-39 derived from GenBank sequences EU290066 and NR_025989 of *M. ovipneumoniae*. Samples with cT values <40 were considered positive. Data are shown as 40 minus the measured cT value to enable semi-quantitative comparison between samples.

### M. ovipneumoniae *serology*

Analysis of serum samples for the presence of *M. ovipneumoniae*-reactive antibodies was performed at WADDL using the laboratory’s monoclonal antibody-based competitive enzyme-linked immunosorbent assay (cELISA) test, which has a diagnostic sensitivity of 88% and a diagnostic specificity of 99.3%. Validation data for the assays may be obtained directly from the laboratory at http://www.vetmed.wsu.edu/depts_waddl/). Data are shown as % inhibition and represent the reduction in binding of the labelled monoclonal antibody to the *M. ovipneumoniae* test antigen caused by competitive binding of serum antibodies from the diagnostic samples. Inhibition of >50% was considered a positive result, and inhibition between 40% and 50% was considered indeterminate.

### Statistical analyses

Data were analyzed by GraphPad Prism, version 9.0 (San Diego, CA) and are shown as mean ± standard deviation (SD). Differences between groups were analyzed by 2-way ANOVA with Sidak’s or Dunnett’s multiple comparisons test or by Student’s *t* test and were considered significant at *P*≤0.05.

## Results

### *Supervised lambing and motherless rearing successfully prevent colonization of domestic lambs with* M. ovipneumoniae *and Pasteurellaceae*

Specific-pathogen-free (SPF) lambs were derived by supervised lambing and artificial rearing from a domestic sheep flock with a history of *M. ovipneumoniae* infection. Pathogen exposure of the ewes was determined by serological analysis prior to lambing. The ewes were free from *C. burnetii* and *M. avium ssp. paratuberculosis*, but had variable serological responses to *M. ovipneumoniae* and parainfluenza virus (PI-3) (**Table 3**). Notably, nasal swabs collected at the same time as the serum samples tested negative for *M. ovipneumoniae* by PCR, indicating previous exposure of the sheep, but no active pathogen shedding.

All thirty lambs born from our ewe flock including the eight experimental lambs selected for our study were free from *M. ovipneumoniae* on days 0 and 30 after birth and one week prior to the experimental inoculation (**Table 4**). We also confirmed the absence of upper respiratory tract colonization by Pasteurellaceae, which include the facultative respiratory pathogens *Mannheimia haemolytica* and *Bibersteinia trehalosi*, in all eight experimental lambs at 3 months of age.

### *Application of nasal wash fluids from* M. ovipneumoniae*-infected lambs leads to successful colonization of specific-pathogen free lambs and induces* M. ovipneumoniae-*specific serum antibodies*

Experimental infection of SPF-lambs aged 103-109 days (3-4 months) with *M. ovipneumoniae* was performed by inoculating four SPF lambs with nasal wash fluids from *M. ovipneumoniae* carriers, while a control group was inoculated with PBS. To achieve a monoinfection with *M. ovipneumoniae*, nasal washes were treated with ceftiofur before experimental inoculation. Microbiological analyses of the nasal washes demonstrated that this treatment successfully eliminated *Bibersteinia trehalosi* and *Mannheimia haemolytica*, which were present at low to moderate levels in the nasal washes prior to antibiotic treatment (data not shown). *M. ovipneumoniae* was detected in pooled nasal wash fluid both before and after ceftiofur treatment. Following inoculation, lambs were monitored for clinical symptoms for twelve weeks, and nasal swabs, serum and BALs for laboratory analyses were collected as shown in **Figure 1A**. All lambs in the *M. ovipneumoniae* group, but none of the lambs in the control group, showed positive PCR results for *M. ovipneumoniae* at two weeks post infection (p.i., **Fig. 1B**). *M. ovipneumoniae* levels peaked at 2–4 weeks, declined at 6 weeks and then plateaued. Colonization with *M. ovipneumoniae* was confirmed by successful culture of viable mycoplasma in nasal swab samples from two of the experimental lambs at 4 weeks post inoculation. We did not detect Pasteurellaceae in any nasal swab samples collected on days 0, 56, and 84 of the infection experiment using standard microbiology techniques (**Table 4**).

**Figure 1:**
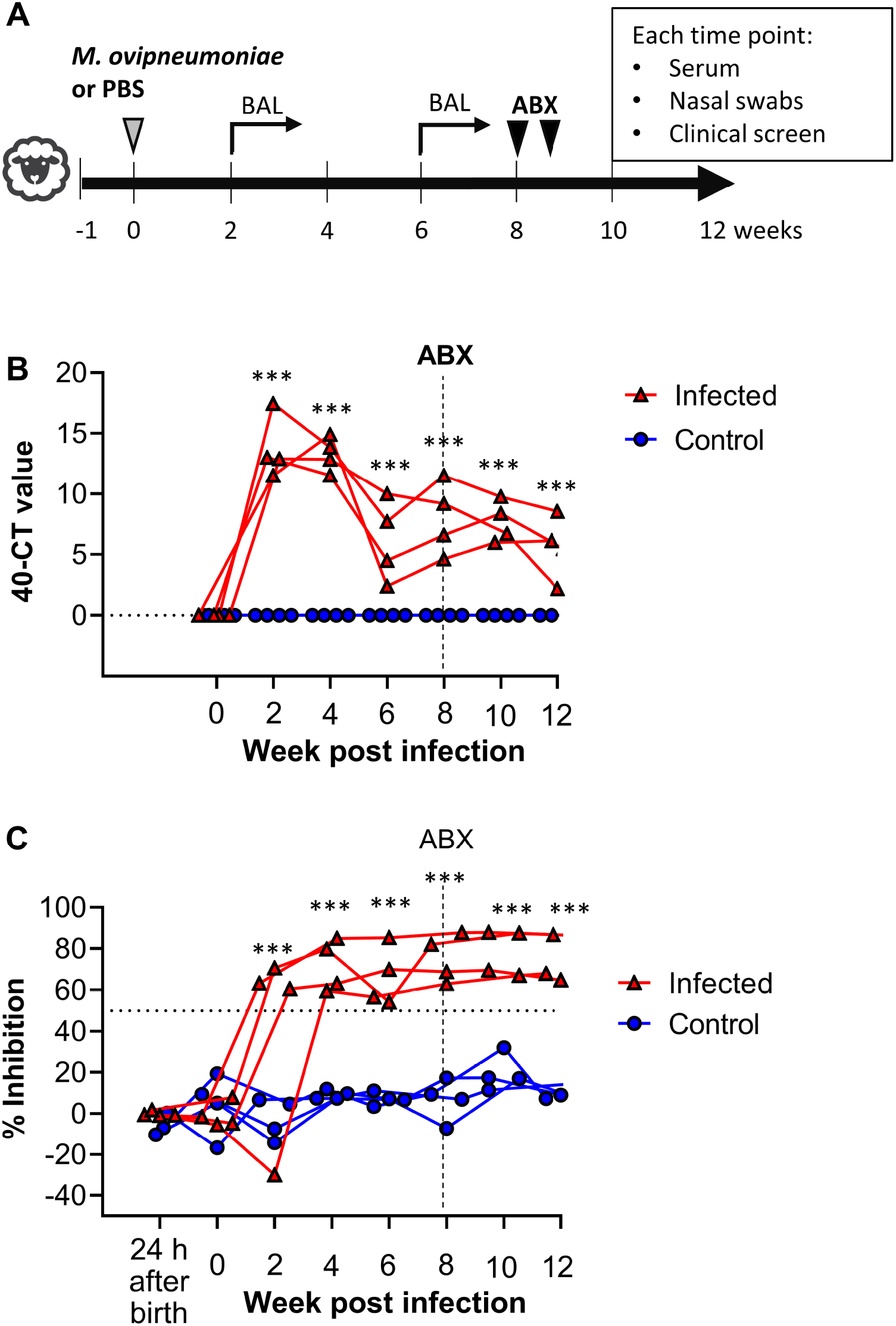
*M. ovipneumoniae* colonization and antibody responses in experimentally infected lambs. **(A)** Experimental schedule. Four SPF lambs aged 3 – 4 months were inoculated with *M. ovipneumoniae*-positive nasal washes, and four control lambs were mock-infected with PBS. All lambs received an antibiotic treatment (gamithromycin) after 8 weeks and were monitored for a total of 12 weeks. **(B)** *M. ovipneumoniae* infection levels in nasal swab samples were determined by qPCR. Data are shown as 40 minus cT value. Triangles represent individual lambs from *M. ovipneumoniae* infection group, circles represent lambs from control group. Data were analyzed by 2-way ANOVA with Dunnett’s multiple comparisons test for differences between week 0 and other time points. \*\*\**P*≤0.001 for the *M. ovipneumoniae-*infected lambs. (**C**) *M. ovipneumoniae-*specific antibody levels were determined by competitive ELISA analysis of serum samples. Red triangles represent individual lambs from *M. ovipneumoniae* infection group, blue circles represent lambs from control group. Data were analyzed by 2-way ANOVA with Dunnett’s multiple comparisons test for differences between week 0 and other time points. \*\*\**P*≤0.001 for the *M. ovipneumoniae-*infected lambs.

All infected lambs developed a strong *M. ovipneumoniae-*specific antibody response that peaked at 4 weeks p.i. and remained high throughout the experimental period (**Fig. 1C**), whereas no significant *M. ovipneumoniae-*reactive antibodies were detected in the control group. Likewise, no *M. ovipneumoniae-reactive* antibodies were present in any of the animals one day after birth, demonstrating that the colostrum replacer used did not contain any crossreactive antibodies that might have contributed to *M. ovipneumoniae-*resistance through passive antibody transfer.

*M. ovipneumoniae* colonization levels detected at different time points throughout the experiment in the experimentally infected lambs by qPCR resembled those seen in lambs of similar ages that were naturally infected with *M. ovipneumoniae* and that served as donor lambs for the nasal wash inocula (**Fig. 2A**). Likewise, antibody levels detected by ELISA in the experimentally infected lambs at multiple time points did not differ significantly from antibody levels detected in the ewes (**Fig. 2B**). These observations suggest that our experimental *M. ovipneumoniae* infection closely replicated natural infection with regards to pathogen load and immune response.

**Figure 2:**
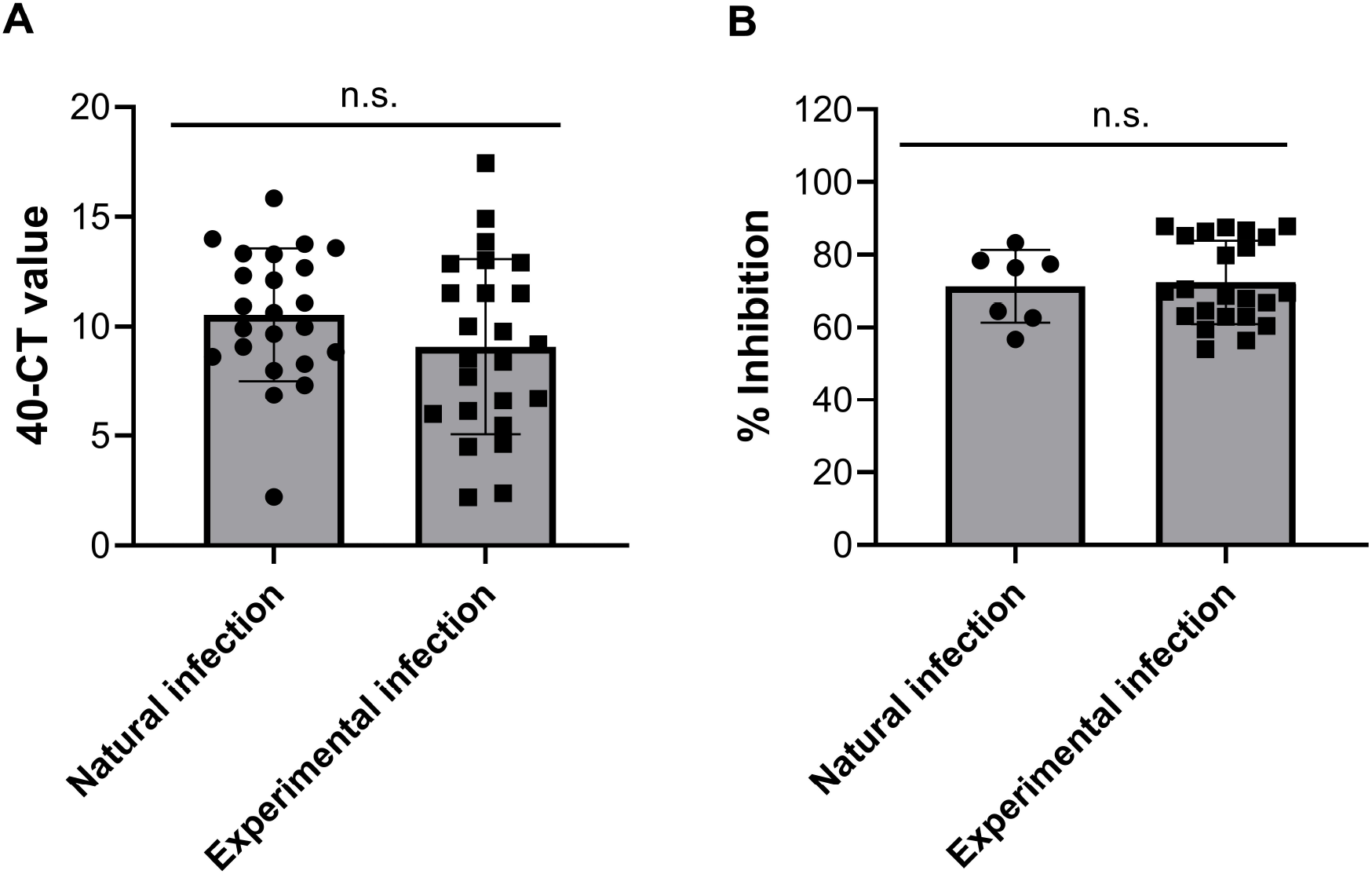
Comparison of *M. ovipneumoniae* levels and serum antibody response to *M. ovipneumoniae* in experimentally and naturally infected domestic sheep. **(A)** *M. ovipneumoniae* infection levels in nasal swab samples were determined by qPCR. Naturally infected animals were lambs of similar ages that tested positive for *M. ovipneumoniae* and that served as donors for the nasal washes. Experimentally infected lambs were the SPF lambs shown in Fig. 1B; symbols represent pooled individual samples from multiple different lambs and time points. Differences between groups were analyzed by Student’s *t* test. **(B)** Antibody levels in the ewes (mothers) compared to the experimentally infected lambs from this study. Only data from animals with positive detection were included. Data from experimental lambs represent multiple animals and time points. Differences between groups were analyzed by Student’s *t* test.

### M. ovipneumoniae *monoinfection in domestic SPF lambs does not cause clinical disease*

Following experimental inoculation, all lambs were closely monitored for symptoms of respiratory disease by clinical scoring (**Table 1**). Interestingly, no significant increase in clinical symptoms was seen in the *M. ovipneumoniae*-infected compared to the uninfected lambs throughout the 12-week experiment (**Fig. 3A-F**). While increased respiratory rates, panting and labored breathing were detected in some of the *M. ovipneumoniae-inoculated* lambs two to three weeks after infection, these observations did not reach significance (**Fig. 3E**). Appetite and body temperatures varied widely in both groups, and significantly decreased appetite was seen in the control group at several time points (**Fig. 3C,D**). As described in more detail below (Fig. 6), lambs in both groups had increased clinical scores and received antiinflammatory treatment following administration of the antibiotic gamithromycin (**Fig. 3A,F**). Lamb body weights and daily gains measured between days 20 and 55 of the experiment, prior to administration of antibiotics, also did not differ significantly between *M. ovipneumoniae*-infected and control lambs (**Fig. 4A, B**). These data indicate that *M. ovipneumoniae* alone did not cause respiratory symptoms in domestic SPF-lambs under controlled laboratory conditions.

**Figure 3:**
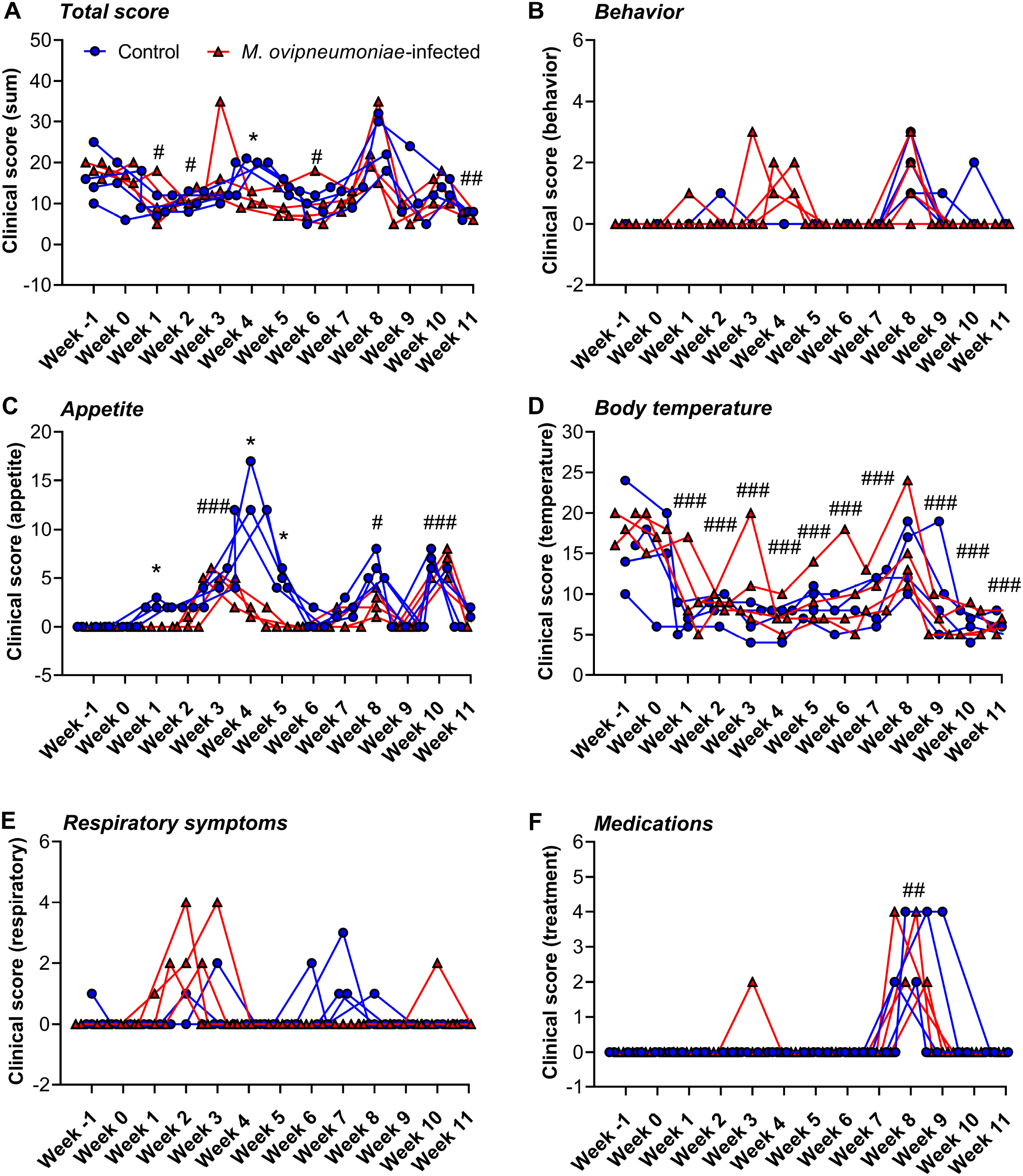
SPF lambs with *M. ovipneumoniae* monoinfection do not develop clinical symptoms of respiratory disease. All lambs were screened twice daily for changes in behavior and appetite and for symptoms of respiratory disease; body temperatures and administered medications were also recorded. Scores were determined using the criteria listed in **Table 1**. Weekly scores are sums of all 14 scores obtained for each lamb per week and per category. Red triangles represent individual lambs from *M. ovipneumoniae* infection group, blue circles represent lambs from control group. (**A**) Total clinical scores, sum of B-F; (**B**) Behavior scores, (**C**) Appetite scores, (**D**) Body temperature scores; (**E**) Respiratory score; (**F**) Medication score. Data were analyzed by 2-way ANOVA with Sidak’s multiple comparisons test for differences between the *M. ovipneumoniae*-infected and control groups. \**P*≤0.05; \*\**P*≤0.01; \*\*\**P*≤0.001. Differences between individual time points and baseline values (week −1 p.i.) for all lambs are indicated by #*P≤*0.05; ##*P*≤0.01; ###*P*≤0.001.

**Figure 4:**
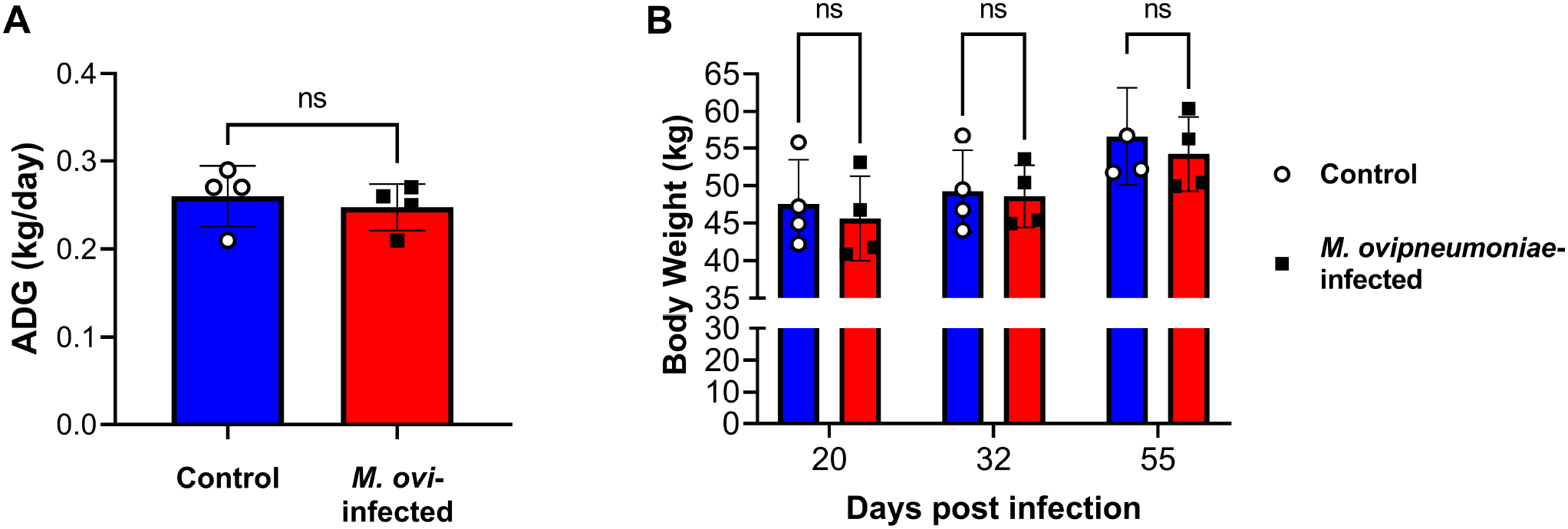
*M. ovipneumoniae* infection does not alter lamb body weights or average daily gains. (**A**) Average daily gains in *M. ovipneumoniae*-infected and control lambs between days 20 and 55 post infection. Differences between groups were analyzed by Student’s *t* test. (**B**). Lamb body weights on days 20, 32 and 55 post infection. Data were analyzed by 2-way ANOVA with Sidak’s multiple comparisons test for differences between the *M. ovipneumoniae*-infected and control groups.

In addition, we analyzed BAL samples collected at 14 and 42 days post infection for evidence of pulmonary inflammation and damage by performing differential cell counts and a lactate dehydrogenase assay. As shown in **Fig. 5A-C**, >90% of cells present in BAL had a typical alveolar macrophage morphology, with oval or round nuclei and cytoplasmic vacuoles. A small number of lymphocytes, neutrophils and eosinophils were also detected (**Fig. 5C**).

**Figure 5:**
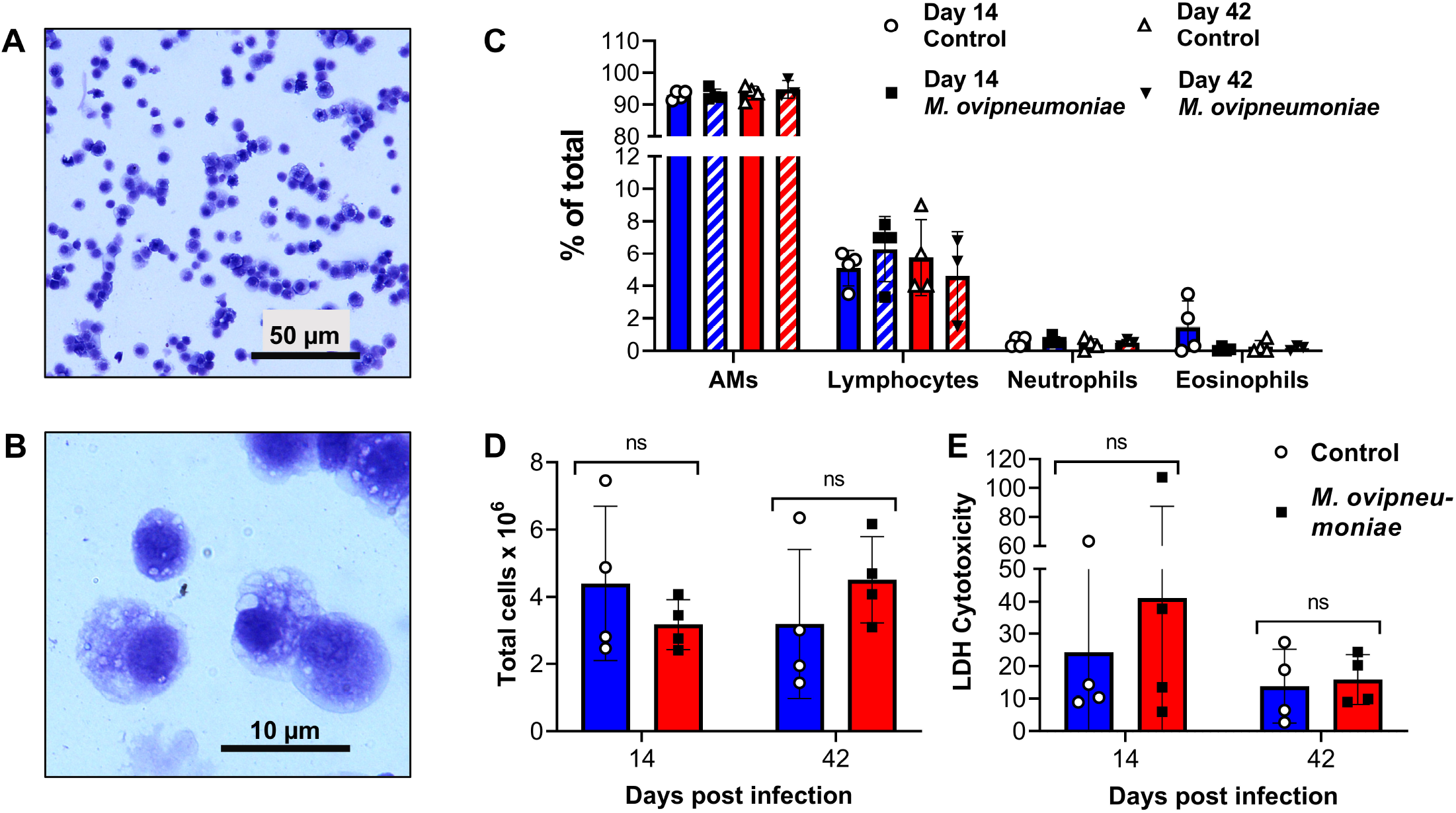
Bronchoalveolar lavage fluid from SPF-lambs experimentally infected with *M. ovipneumoniae* shows no evidence of lung inflammation. **(A, B)** Cellular composition of BAL from an *M. ovipneumoniae*-infected lamb 14 days after experimental inoculation. Representative image from one of four infected sheep shows typical macrophage morphology of the majority of the cells at (A) low and (B) high magnification. **(C)** Differential cell counts of BAL collected on days 14 and 42 after experimental inoculation from *M. ovipneumoniae*-infected and non-infected lambs (n=4). **(D)** Total cell counts of BAL collected on days 14 and 42 after experimental inoculation from *M. ovipneumoniae*-infected and mock-infected lambs (n=4). **(E)** Analysis of lactate dehydrogenase in cell-free supernatants of BAL collected on days 14 and 42 post inoculation. Individual data points and mean ± SD are shown. Differences between groups were analyzed by Student’s *t* test.

Neither cell composition nor total cell counts differed significantly between the *M. ovipneumoniae-*infected and non-infected lambs (**Fig. 5C, D**). We also analyzed the cell-free supernatants for the presence of lactate dehydrogenase (LDH) as a marker of lung injury [30]. Although there was a trend for increased LDH release upon *M. ovipneumoniae* infection 14 days after experimental inoculation, no significant differences between the two groups were found at either of the time points analyzed (**Fig. 5E**). These data suggest that the experimental infection did not cause pneumonia in the SPF lambs, consistent with the clinical findings presented above.

### *High dose treatment with the macrolide antibiotic gamithromycin fails to eliminate* M. ovipneumoniae *colonization*

We next sought to determine whether subclinical *M. ovipneumoniae* infection could be eliminated by antibiotic treatment with gamithromycin (Zactran®), which is currently used therapeutically to treat *M. bovis* infection in cattle and *M. hyopneumoniae* infection in swine [23, 24]. Eight weeks after initiation of experimental *M. ovipneumoniae* infection, all lambs were treated with two doses of gamithromycin on days 56 and 61 post infection (**Fig. 6A**). However, the antibiotic treatment neither eliminated nor significantly decreased the level of *M. ovipneumoniae* infection as determined by qPCR in any of the lambs either at 2 or at 4 weeks after the treatment (**Fig. 6B**). Notably, significant side effects of the gamithromycin treatment including changes in behavior, decreased appetite, and increased body temperatures, were observed after treatment in both the *M. ovipneumoniae*-infected lambs and the control group (**Fig. 6C**). To counteract these side effects, anti-inflammatory treatment was administered to some of the animals (**Fig. 3F**). These data strongly indicate that gamithromycin is unsuitable for treatment of *M. ovipneumoniae* infection in domestic lambs, since it does not eliminate infection, but has considerable adverse effects.

**Figure 6:**
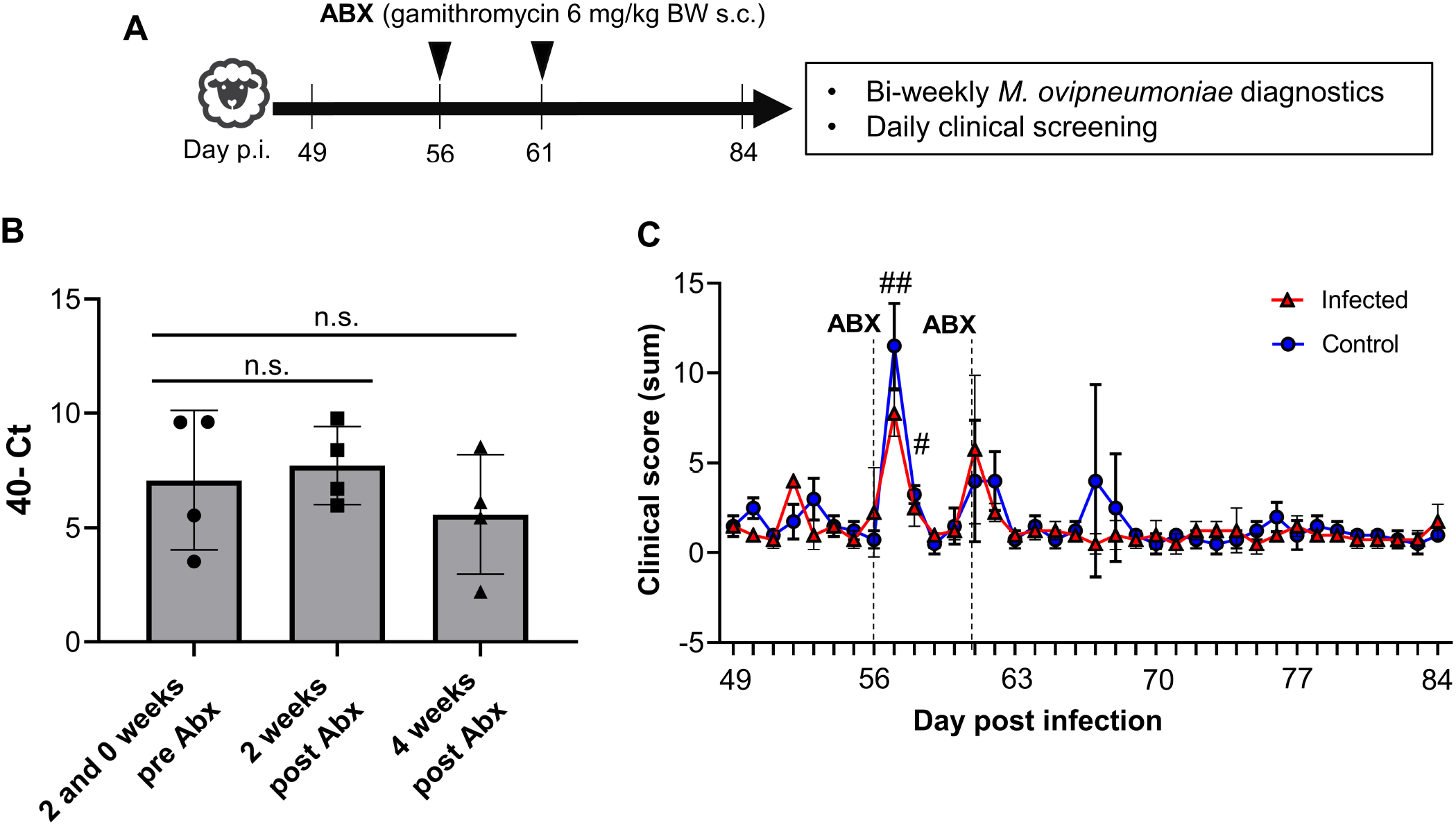
Antibiotic treatment with gamithromycin does not eliminate asymptomatic *M. ovipneumoniae* infection. **(A)** Detailed timeline for antibiotic administration. (**B**) *M. ovipneumoniae* infection levels in nasal swab samples of lambs inoculated with *M. ovipneumoniae-*positive nasal washes were determined by qPCR at 14 and 0 days before administration of gamithromycin (average of two time points) and at 14 and 28 days post administration of gamithromycin. Data are shown as 40 minus cT value. (**C**) Clinical scores of *M. ovipneumoniae*-positive and negative lambs after gamithromycin treatment show side effects of antibiotic administration. All lambs were screened twice daily for changes in behavior and appetite and for symptoms of respiratory disease; body temperatures and administered medications were also recorded. Scores were determined using the criteria listed in Table 1. Daily scores reflect the sum of two scores per day for all five criteria. Red triangles represent individual lambs from *M. ovipneumoniae* infection group, blue circles represent lambs from control group. Significant differences (*P*≤0.05) between individual time points and baseline values (day 49 p.i.) for all lambs are indicated by #.

## Discussion

*Mycoplasma ovipneumoniae* is a key co-factor in the pathogenesis of chronic, atypical pneumonia in sheep, which has been associated with production losses in domestic lambs and significant deaths in wild sheep populations. In our study, we sought to define whether *M. ovipneumoniae* monoinfection causes clinical respiratory disease in immunologically naïve, specific-pathogen-free domestic lambs. We also tested whether established *M. ovipneumoniae* infection could be eliminated by antibiotic treatment. We found that *M. ovipneumoniae* caused stable, asymptomatic colonization of the upper respiratory tract that induced a significant antibody response, but that could not be cleared by gamithromycin application.

Whether *M. ovipneumoniae* causes respiratory disease in healthy domestic lambs has been a matter of debate [16–19, 31]. Our tightly controlled experimental conditions ensured that experimental lambs were not exposed to respiratory pathogens other than the *M. ovipneumoniae* used in the infections, leading to true *M. ovipneumoniae* monoinfections. The twice-daily clinical screens and cellular and molecular analyses of the bronchoalveolar lavage fluid demonstrated that active *M. ovipneumoniae* colonization of the upper airways did not lead to clinical disease or lung pathology in our model. In contrast to recent studies, we also did not observe any differences in lamb body weights or daily gains associated with *M. ovipneumoniae* infection [6, 7]. Multiple different hypotheses have been proposed to explain why *M. ovipneumoniae* causes asymptomatic airway colonization in some instances, but severe pneumonia in other cases: (1) differences in *M. ovipneumoniae* strain virulence; (2) age-related differences in disease susceptibility; (3) differences in the immune response to *M. ovipneumoniae;* and (4) presence of additional facultative pathogens such as *Mannheimia haemolytica*.

Asymptomatic *M. ovipneumoniae* infections as seen in our study may be caused by experimental inoculation with non-pathogenic *M. ovipneumoniae* variants, since *M. ovipneumoniae* strains are highly diverse and can differ in their virulence [32]. Moreover, *Mycoplasma* may lose their virulence upon prolonged culture [32]. Such unintended, culture-dependent attenuation of *M. ovipneumoniae* isolates has been discussed as the cause for asymptomatic *M. ovipneumoniae* infection in several previous studies [19, 20]. However, the inocula used in our study were fresh nasal washes obtained from *M. ovipneumoniae*-positive lambs that showed symptoms of respiratory disease, strongly suggesting that the lack of clinical symptoms upon experimental *M. ovipneumoniae* infection were not due to the use of an avirulent or attenuated *M. ovipneumoniae* strain.

With regard to the second hypothesis, it is generally accepted that clinical infections with *M. ovipneumoniae* most commonly occur in lambs under one year of age, while adult sheep serve as asymptomatic *M. ovipneumoniae* carriers [3, 33]. Plowright et al. showed that, in bighorn sheep, prevalence of *M. ovipneumoniae* was highest in lambs and aged animals, but low in adults, pointing to age-dependent infection dynamics [34]. However, it is unclear whether the increased susceptibility of lambs was due to the lack of established adaptive immune responses to *M. ovipneumoniae* or other age-related differences in respiratory anatomy, physiology, or immunity. Interestingly, Gilmour et al. found increased lung pathology in seven-month-old lambs compared to five-week-old lambs upon experimental *M. ovipneumoniae* infection [31]. In bighorn sheep, yearlings and older adults with no previous *M. ovipneumoniae* infection were equally susceptible to *M. ovipneumoniae-*induced pneumonia [20]. Thus, age-dependent mechanisms alone do not explain the variable susceptibility of sheep to *M. ovipneumoniae* disease.

In bighorn sheep, introduction of *M. ovipneumoniae* into herds with no prior exposure to the pathogen leads to particularly devastating disease outbreaks, suggesting that immunologically naïve populations are especially prone to symptomatic infections. *In vitro* studies have shown that *M. ovipneumoniae* specific antibodies can have protective functions by mediating opsonization and phagocytosis of the Mycoplasma [35]. Likewise, Niang et al. [36] found increased levels of *M. ovipneumoniae-reactive* antibodies in lambs that had recovered from the clinical *M. ovipneumoniae* infection. In our study, lambs had no detectable *M. ovipneumoniae* specific antibodies prior to experimental inoculation but developed a significant, stable antibody response within two weeks after infection, confirming previous studies that have tracked humoral responses to *M. ovipneumoniae* [37]. These observations indicate that lack of existing immunity does not lead to more severe disease in *M. ovipneumoniae* infected sheep. Interestingly, the development of an antibody response was not associated with decreased *M. ovipneumoniae* colonization in our study. A possible explanation for this phenomenon is that antibody levels measured in serum consist mainly of IgG, which may have protective functions within the lungs, whereas a strong mucosal IgA response may be necessary to eliminate asymptomatic *M. ovipneumoniae* colonization from the upper respiratory tract.

Overall, our findings that *M. ovipneumoniae* application in domestic lambs did not cause respiratory disease were in line with earlier studies that showed that pneumonia develops as a result of *M. ovipneumoniae* co-infections with other facultative bacterial pathogens [9, 15, 19, 38]. In bighorn sheep, where a large number of studies have been performed, *M. ovipneumoniae* is likely essential for causing severe respiratory disease [39], but consistent clinical disease and lung pathology only developed when additional agents such as *Mannheimia haemolytica, Bibersteinia trehalosi, Pasteurella multocida* or *Fusobacterium necrophorum* [9, 13, 38] also were present. In further support of this hypothesis, the lambs that served as donors for the nasal wash fluids that we used to inoculate the experimental animals in our study showed symptoms of respiratory disease and tested positive for *Mannheimia haemolytica* and *Bibersteinia trehalosi* in addition to the *M. ovipneumoniae*.

Even if *M. ovipneumoniae* alone does not cause clinical respiratory disease in domestic sheep, an effective treatment strategy that eliminates *M. ovipneumoniae* infection from domestic sheep flocks could reduce production losses due to complex respiratory disease and protect bighorn sheep from devastating pneumonia epizootics. To address this issue, we analyzed whether antibiotic therapy could eliminate *M. ovipneumoniae* colonization in our asymptomatically infected lambs. We chose the macrolide antibiotic gamithromycin (Zactran®), because gamithromycin application in goats infected with *Mycoplasma* and *Mannheimia* was successful in treatment of clinical pneumonia [8], and preventive treatment with gamithromycin in cattle was successful for reducing bovine respiratory disease, which commonly involves *Mycoplasma spp*. [9]. A recent study by Jay et al. on the antibiotic susceptibility of *M. ovipneumoniae* have showed homogenously low minimum inhibitory concentrations for a wide range of antimicrobials including macrolides, with no indication of significant antimicrobial resistance [4]. Since gamithromycin is currently not approved for use in sheep in the US, the presence of resistant *M. ovipneumoniae* strains in the infected donor sheep was unlikely. However, our data showed no significant reduction in *M. ovipneumoniae* colonization levels in our experimental lambs following two injections of the recommended gamithromycin dose. It remains to be investigated whether failure of the antibiotic treatment was due to bacterial biofilm formation [40], intracellular location of the mycoplasma [41] or an alternative antimicrobial resistance mechanism. Notably, all lambs developed significant side effects following gamithromycin administration, including increased body temperatures, decreased appetites and altered behavior as well as local inflammation at the injection sites that warranted the application of analgesics. These side effects were greatly more severe than the transient mild to moderate swellings at the site of injection described for this drug by the European Medicines Agency [42].

Why domestic lambs tolerate *M. ovipneumoniae* infection, while bighorn lambs suffer severe clinical disease may thus be due to host genetics and will require further investigations. The inoculation approach that involved nasal, oral and ocular application of 4-months-old domestic SPF lambs with ceftiofur-treated nasal washes from naturally infected, symptomatic lambs led to 100% successful upper airway colonization with *M. ovipneumoniae*, but not with Pasteurellaceae, confirming earlier studies by Thomas Besser’s group [20, 27, 28]. Colonization levels as determined based on the cT values in the qPCR reactions was similar in experimentally and naturally infected lambs, and antibody levels measured by *M. ovipneumoniae-specific* ELISA also closely replicated antibody levels seen upon natural infection. Thus, our infection model represents a valid tool to investigate the role of *M. ovipneumoniae* in sheep respiratory disease. In future studies, we plan to utilize this *M. ovipneumoniae* infection model to explore novel preventative or treatment approaches and to define how interactions between *M. ovipneumoniae* and other facultative respiratory pathogens lead to clinical respiratory disease and pulmonary pathology in domestic sheep.

## Acknowledgements

We gratefully acknowledge support by NIFA AFRI Animal Health & Disease Program grant no. # 2018-06895 from the USDA National Institute of Food and Agriculture for this work. This study was also supported by the Montana Agricultural Experiment Station, USDA/NIFA Animal Health funds and the Johnson Family Foundation. Our sincerest thanks go to the sheep team at the Johnson Family Livestock Facility and to Dr. Bruce Sorensen for providing excellent care of our animals. We also would like to thank Andy Sebrell, Marziah Hashimi and Farimah Moghimpour for assisting with sample processing throughout our study. We also thank Dr. Patrick Hatfield, MSU Department of Animal and Range Sciences, for providing access to lambs naturally infected with *M. ovipneumoniae*. Many thanks go to Dr. Martin Ganter, Veterinary University of Hannover, Germany, for helpful discussions during the planning phase of this study. Finally, we would like to thank Daniel Bradway and all the other staff from the Washington Animal Disease Diagnostic Laboratory for sample analysis.

## Author Contributions

DB, MJ, ARA, TJ, KJ and TB designed the study; TJ, KJ, BTJ and JS performed the experiments; TJ, BTJ and DB analyzed the data; DB, TJ, ARA and MJ interpreted the data; DB obtained funding for the study; DB and TJ wrote the manuscript; all authors have read and approved the final manuscript.

